# Self-organization of PIP3 signaling is controlled by the confinement, topology, and curvature of the cell membrane

**DOI:** 10.1101/2024.01.20.576405

**Authors:** Sema Erisis, Marcel Hörning

## Abstract

PIP3 is a signaling lipid on the plasma membrane that plays a fundamental role in cell signaling with a strong impact on cell physiology and diseases. It is responsible for the protruding edge formation, cell polarization, macropinocytosis and other membrane remodeling dynamics in cells. It has been shown that the membrane confinement and curvature affects the wave formation of PIP3 and Factin. But even in the absence of F-actin, a complex self-organization of the spatiotemporal PIP3 waves is observed. In recent findings, we have shown that these waves can be guided and pinned on strongly bended *Dictyostelium* membranes caused by molecular crowding and curvature limited diffusion. Based on these experimental findings, we investigate the spatiotemporal PIP3 wave dynamics on realist 3D cell-like membranes to explore the effect of curvature limited diffusion, as observed experimentally. We use an established stochastic reaction-diffusion model with enzymatic MichaelisMenten type reactions that mimics the dynamics of *Dictyostelium* cells. As these cells mimic the 3D shape and size observed experimentally, we found that the PIP3 wave directionality can be explained by a Hopf-like and a reverse periodic-doubling bifurcation for uniform diffusion and curvature limited diffusion properties. Finally, we compare the results with recent experimental findings and discuss the discrepancy between the biological and numerical results.

## Introduction

Many features of pathways observed in the model organism *Dictyostelium discoideum* are similar to those in eukaryotic cells (Martín-González et al., 2020), such as directional migration, cell adhesion, gene expression, and cell-cell signaling (Manahan et al., 2004). In both organisms, the Phosphatidylinositol (3,4,5)-trisphosphate (PIP3) molecule acts as a pivotal player in orchestrating cellular responses, serving as a key regulator for directional migration and cell polarization (Arai et al., 2010; Posor et al., 2022). The self-organized patterns of PIP3 waves observed in *Dictyostelium* cells have drawn attention for their relevance to similar processes in mammalian cells, and have been used as an alternative experimental model system for many human diseases (Williams, 2010; Mathavarajah et al., 2017; Haver and Scaglione, 2021; Storey et al., 2022).

PIP3 is a triply phosphorylated membrane lipid involved in the signaling pathway to recruit F-actin (Beta et al., 2023). It is generated from phosphatidylinositol 4,5-bisphosphate (PIP2) by adding a phosphate group through phosphoinositide 3-kinase (PI3K), and then converted back by the phosphatase and tensin homolog (PTEN) (Kölsch et al., 2008). While PIP3 is correlated to other signaling activities, such as to Rac (Innocenti et al., 2003; Graziano et al., 2017), it leads to activation of the mTOR signaling pathway, and hence, plays an important role in the cytoskeleton organization (Tariq and Luikart, 2021). Nano-topological induced changes to the membrane of *Dictyostelium* cells trigger a pronounced rearrangement of the F-actin and PIP3 wave patterns (Honda et al., 2021; Yang et al., 2023). No F-actin wave patterns are observed upon Latrunculin A treatment (Coué et al., 1987). However, even in the absence of F-actin, local membrane curvature pins and guides the PIP3 waves locally (Hörning et al., 2021). The size and shape of the cell dictate the PIP3 wave dynamics on the membrane. Smaller cells lead to transient spot waves and larger ones to rotating waves (Hörning and Shibata, 2019). Only certain unstable modes with a wavelength limited by the cell size can be realized, as the size of the cell dictates the dynamics trough the excitable dispersion and restitution properties of the cell (Burkart et al., 2022).

The confinement of the membrane, i.e. finite size effect, has been experimentally observed in the PAR cell polarity network of *C. elegans*, where a decrease in the cell size of the embryos destabilizes polarity and induces premature loss of division asymmetry (Hubatsch et al., 2019). That is because the volume of a sphere increases faster than its surface area, as the volume scales with the cell radius *R* as *R*^3^, whereas the cell surface scales as *R*^2^. Dynamics on the membrane have been studied also from the theoretical and general biophysical point of view to explain biological systems, as cell membranes are active deformable surfaces (Miller et al., 2018; Haas and Goldstein, 2021; Nishide and Ishihara, 2022; Yin et al., 2022; Hirashima and Matsuda, 2024). In these studies, the curvature of the membrane plays an important role to describe the self-organization of signals and membrane deformation in various biological systems, as the additional effect of curvature on the diffusion increases the complexity to understand and predict wave dynamics (Gov, 2006; Frank et al., 2019).

Also in *Dictyostelium* cells, membrane curvature plays an essential role for the guidance of cell migration. In curved nano-topological environments, it has been shown that the membrane curvature affects the wave formation of PIP3 and F-actin (Honda et al., 2021; Yang et al., 2023). That means that the cells actively sense and react to physical cues of the environment. The results of Yang et al. (Yang et al., 2023) indicate that the nanotopography is sensed directly by the cytoskeletal excitable network, whereas the signal-transduction excitable network is only indirectly affected due to feedback dynamics between both networks. Contrarily, we have shown that membrane curvature alone affects the self-organization of the PIP3 waves even in the absence of the cytoskeletal excitable network by the analysis of three-dimensional actin-polymerization inhibited *Dictyostelium* cell membranes(Hörning et al., 2021). This is explained by the combination of molecular crowding and membrane curvature limited diffusion, which means that PIP3 waves are guided or even pinned as spiral waves depending on the strength of membrane curvature. The additional effect of the membrane confinement and topology leads to an pronounced PIP3 wave directionality along the equatorial plane of the cell (Hörning and Shibata, 2019).

In this study, we numerically investigate the difference of the self-organization of spatiotemporal PIP3 waves on actin-polymerization inhibited cells with and without curvature limited diffusion on the membrane, to verify the PIP3 dynamics observed experimentally, and gain insights of the fundamental nature of PIP3 pattern formation in the absence of the cytoskeletal network. We use a stochastic reaction diffusion system with enzymatic Michaelis-Menten type reactions on a three-dimensional membrane, that closely resembles actin-polymerization inhibited *Dictyostelium* cells. By variations in size and shape of the cells similar to that observed experimentally, we compare and quantify the restitution dynamics and PIP3 wave propagation direction on the membrane. We found a Hopf-like bifurcation for membrane topologies with uniform diffusion, and a reverse period-doubling bifurcation for membranes with curvature limited diffusion at the contact line for intermediate-sized non-spherical cells. That means that either dominant longitudinal or transversal stable PIP3 wave dynamics were observed. Larger cells showed secondary PIP3 wave formations, which led to chaotic wave formations. In the latter part of this work, we compare our results with the experimentally quantified results of *Dictyostelium* cells that partly recover the complex dynamics. Based on these results, we discuss possible scenarios that may explain the similarity and discrepancy between the biological and numerical results.

## Methods

The stochastic model was implemented in C, and the mesh preparation and computational analysis of the computed dynamics in MATLAB (The MathWorks, Natick, MA). A detailed description of the mesh preparation and computational analysis was published before for the analysis of experimentally observed *Dictyostelium* cells (Hörning and Shibata, 2019; Hörning et al., 2021) together with a basic version of the implemented code that is available at MATLAB Central File Exchange (Hörning, 2021).

### Computational Platform

The stochastic model was computed on a NEC cluster using 20 cores (Intel Xeon Gold 6138 with 2.00GHz, Skylake, 92GB memory). The server architecture was provided by the High-Performance Computing Center Stuttgart (HLRS), Germany. A single simulation of 5000 s lipid membrane dynamics took about 24 h of simulation time at a single core.

### Membrane topology and mapping

The four spherical membrane topologies with *r*_A_ between 0% and 30% were generated using a Delaunay triangulation routine (Persson and Strang, 2004). The number of nodes for the four topologies was kept between about 3500 and 4000 nodes to ensure accurate mesh generation. Lists of the spatial position of the nodes and meshes, the index of the meshes, and their neighboring meshes, as well as the boundary length between meshes were stored and provided to the stochastic model.

The meshes were visualized using the Mollweide projection, a homolographic equal-area projection, i.e. the area accuracy-based projection (Weisstein, 2020). The spherical coordinates (*ϕ, θ*) were transformed to cartesian coordinates (*x, y*), as follows:

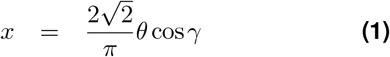

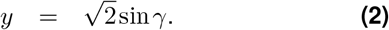

The auxiliary angle *γ* is given as

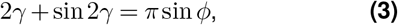

and iteratively solved by the Newton-Raphson method (Weisstein, 2020; Snyder, 1987).

### Tracking of signaling

For tracking the PIP3 lipid domains, spherical harmonic analysis was applied to smoothen the computed intensity distributions *I*_mem_ (*ϕ, θ*) on the membrane, which enabled peek detection. The smoothed intensity distribution *Ĩ*mem (*ϕ, θ*) was computed up to the scalar harmonic degree *l* = 3 (octupole moment) for each time step, as follows

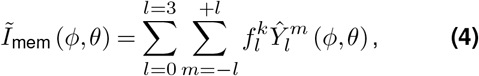

where the spherical harmonics is given by

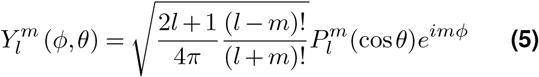

with the Legendre Polynoms 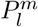 and 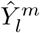 the complex conjugated of 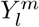. The spherical harmonic coefficients are defined as

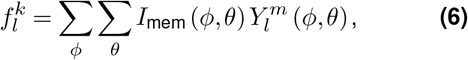

where *ϕ* and *θ* denote the discrete and isotropically distributed node positions of the mesh.

The velocity and the angular velocity components of the tracked domains were calculated as

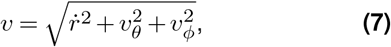

where the angular velocity components are

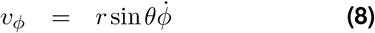

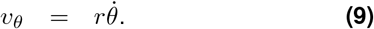

## Results

### Self-organized Lipid Signaling in Membranes

Phosphatidylinositol lipid dynamics have been modeled before on the basis of experimental observations of *Dictyostelium* cells (Arai et al., 2010). A simple reaction diffusion system has been introduced that captures the spontaneous phosphatidylinositol lipid dynamics on a one-dimensional ring, resembling the cell membrane periphery. Based on that model an improved reaction diffusion system has been introduced with enzymatic Michaelis-Menten type reactions (Shibata et al., 2012). In that model, the reactions between PIP2 and PIP3 are not isolated, and the total concentrations of both lipids can change with time independently of PI3K and PTEN. The reaction scheme is given as

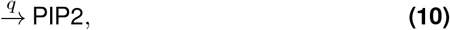

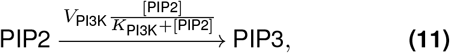

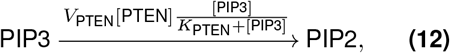

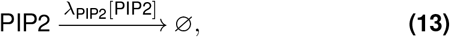

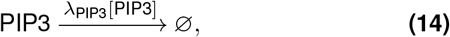

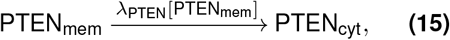

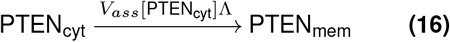

with

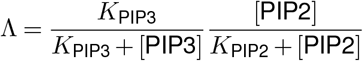

and

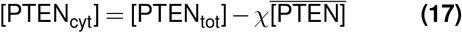

where [PIP2], [PIP3], and [PTEN] are the membrane concentrations of PIP2, PIP3 and PTEN, respectively, and *q, χ, V*_i_, *K*_i_ and *λ*_i_ are reaction constants (see Supplementary Materials Table 1). [PTENcyt], [PTENtot], and 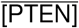 are the cytosolic, total and average membrane concentration of PTEN.

In this study, these reaction diffusion processes are computed on three-dimensional cell shaped surfaces using the *τ* -leaping Gillespie algorithm based method (Gillespie, 2001; Cao et al., 2006). The surfaces resemble actin-polymerization inhibited *Dictyostelium* cells (Hörning and Shibata, 2019), which are generated by a Delaunay triangulation routine (see Section Methods). The diffusion is considered for PIP2 and PIP3 as a stochastic jump process between neighboring grids and the reaction rates (propensity functions on top of the arrows of Eqs. 10-16) with a discrete time stepping of *τ* = 2 × 10−4 s. The dynamics were simulated for a duration of 5 × 103 s for different cell radii ranging from *R* = 5 μm to 16 μm. Additional variations of the membrane topology between *r*_A_ = 0% (spherical) and 30% (flattened) led to 44 different cell topologies. Figure 1A shows the grid of a cell with *r*_A_ = 20%. The contact line between substrate adhered membrane (flat surface) and non-adhered membrane is highlighted by a red line. The diffusion on the membrane surface is considered as *D*_mem_ = 0.2 μm^2^/s equally for PIP2 and PIP3 (Fujiwara et al., 2002). The diffusion at the contact line is considered either as *D*_mem_ = *D*_cl_ or *D*_mem_ ≠ *D*_cl_ = 0.04 μm^2^/s (see Supplementary Materials Fig. S1). We refer from here on to membranes with uniform diffusion and diffusion limited membranes, respectively.

**Figure 1.**
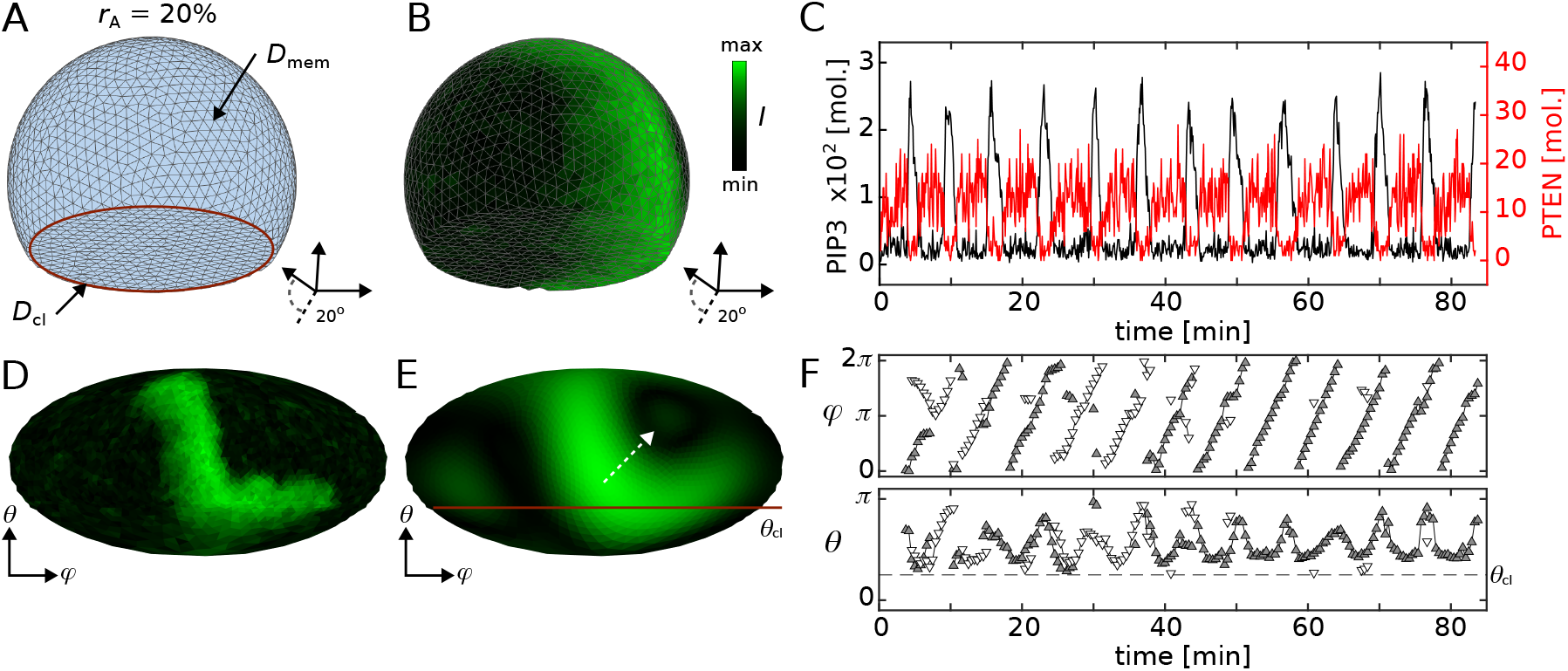
Mesh and signal visualization. **A-B** show the 3D views of a plain mesh and a mesh with mapped computed PIP3 intensity signaling. The examples show meshes with *r*_A_ = 20% from the bottom with an inclination angle of 20*°*. **C** shows the molecular changes over time of PIP3 (black) and PTEN (red) at a single grid. **D-E** show the 2D Mollweide projection of the example shown in **B**, where *θ*_cl_ defines the contact line between adhesive and non-adhesive part of the membrane that is indicated by the red solid line in **A** and **E. E** shows the smoothed intensity distribution obtained by spherical harmonics analysis of the simulated data shown in **B** and **D**. The white dashed arrow indicates the direction of wave propagation from the detected peak. The color bars indicate the normalized PIP3 signaling intensity. **F** shows the spatiotemporal position of the detected wave positions in spherical coordinates. The dashed line indicates the contact line.

Figure 1B shows a snapshot of a typical PIP3 domain distribution on the surface of a cell, and Fig. 1C shows the temporal dynamics of PIP3 (black) and PTEN (red) at a single membrane node. In order to track these PIP3 domains on the membrane over time, spherical harmonics analysis was applied and higher order modes were removed. This simplifies the complex patterns on the surface and enables wave detection. The same approach was applied experimentally at *Dictyostelium* cells before (Hörning and Shibata, 2019). The twodimensional projections (see Section Methods) of that cell are shown in Figs. 1D and E. The unprocessed simulated and the by spherical harmonics simplified PIP3 distributions are shown. The position and direction of the PIP3 domain dynamic are indicated by the white dashed arrow. The horizontal line indicates the contact line, similarly as indicated in Fig. 1A. The area below that line indicates the substrate adhered part of the membrane. Up to two domains at a time were tracked on the surface of the membrane. The spatial position of the waves is tracked by spherical coordinates with the azimuthal angle *φ* (latitude) and the polar angle *θ* (longitude). Figure 1F shows an example. Up to two domains are shown with white and grey triangles at a time. When one PIP3 wave is present (e.g., white), the secondary wave is shown by the opposite color (e.g., grey). Same waves are connected by a black line.

### Membrane Topology and Diffusivity

Depending on the membrane topology various dynamics can be observed (Figs. 2A-D). Perfectly spherical cells (*r*_A_ = 0%) do not have a symmetry axis, so stable rotating waves can be observed in various configurations leading to horizontally and vertically propagating waves. An example of the latter is shown in Fig. 2A. Membrane snapshots show one period of two rotating spiral waves. Their tips, i.e. phase singularities, are located on the opposite side of the cells with a common wave front that travels vertically up and down. This dynamic is also indicated by the vertical lines in the Kymograph, which is taken at the equatorial height of the cell (*θ* = *π/*2). Adherent cells (*r*_A_ *>* 0%), as shown exemplarily in Fig. 2B (*r*_A_ = 20%), have the tendency to exhibit stable vertically rotating waves. The spiral tips are located at the close proximity to the top and the bottom of the cell membrane due to the vertical symmetry axis of the membrane. This is because excitable waves tend to propagate along the longest path on spatially confined membrane surfaces due to their inhibitory signaling properties. In other words, the wave front propagates the path where the excitable PIP3 wave did not occur for the longest time, as the PIP3 wave propagates in its own inhibitory tail. In adherent cells that are the great circles in close proximity to the equatorial line.

**Figure 2.**
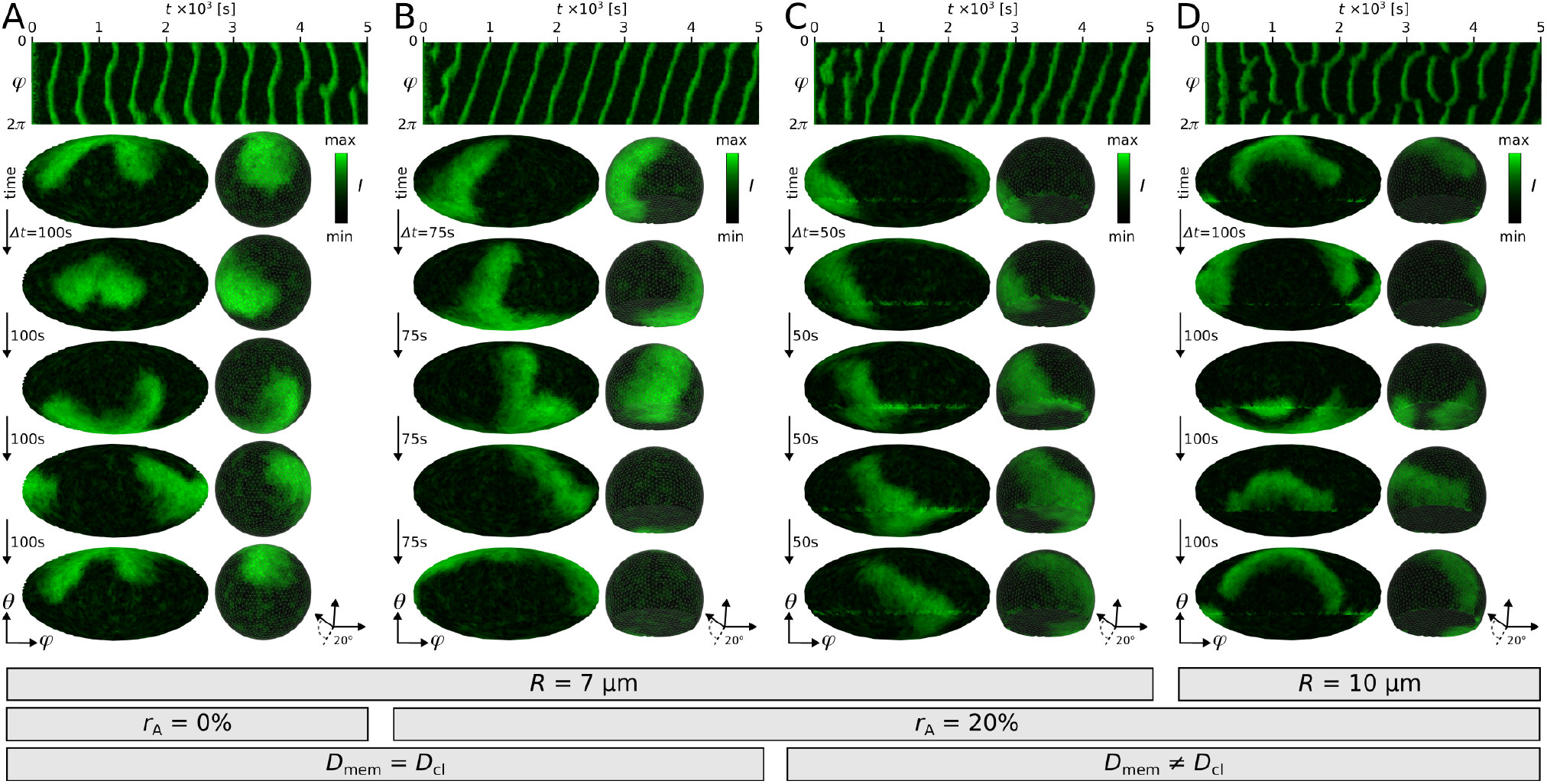
Influence of membrane shape and curvature limited diffusion on PIP3 wave dynamics. **A-D** show typical examples with different membrane properties (*R, r*_A_ and *D*). The left and middle panels are five snapshots 2D map and 3D view of the PIP3 membrane distribution. The right panels show the kymographs obtained at the equatorial section of the cell membrane (*θ* = 0). **A** and **B** show cells with constant diffusion at the entire membrane but different cell shape, **B** and **C** show cells of same shape and geometry but different diffusion properties, and **C** and **D** show cells with curvature limited diffusion at the contact line but different cell size. The 3D views show the ventral membrane from the bottom with an inclination angle of 20*°*. The color bars indicate the normalized PIP3 signaling intensity.

When introducing diffusion limited PIP3 signaling at the contact line of the membrane (*D*_cl_ = 0.04 μm^2^/s), that more closely relates to the experimentally observed actin-polymerization inhibited (Hörning et al., 2021) and motile *Dictyostelium* cells (Honda et al., 2021), stable horizontally rotating waves are observed (Fig. 2C), too. However, no cases of vertically rotating waves are observed, contrarily to cells with uniform diffusibility (*D*_mem_ = *D*_cl_) where also cases of stable vertically rotating waves are observed. Thus, despite the same cell topology (*r*_A_ = 20%, *R* = 7 μm) the diffusion limitation at the contact line leads to the stabilization of the PIP3 wave directionality. In larger cells, more complex PIP3 wave dynamics are observed. Figure 2D shows a *R* = 10 μm sized cell, where the strong horizontal directionality is broken, as a secondary PIP3 wave appears. This is possible, as there is a natural speed limit defined by the dispersion properties of the system, i.e. relationship between wave speed and rotational frequency. The wave needs more time to propagate around the membrane with the increase in cell size, which in turn enables the formation of a secondary PIP3 domain, if the cell is sufficiently large. Figure 3A depicts a simplified one-dimensional scheme for three different radii. The larger the radius of the one-dimensional circle, the larger the wave speed. The wave speed saturates at a certain radius *R*, here *R*_2_ *< R < R*_3_, and leads to longer delays within the wave front and inhibitory wave tail, where the formation of secondary waves may occur. In case of diffusion limited membranes, these secondary waves appear more prominently on the contact line (Fig. 2D, third snapshot), contrarily to membranes with uniform diffusibility, where those waves do not appear at any specific membrane location.

**Figure 3.**
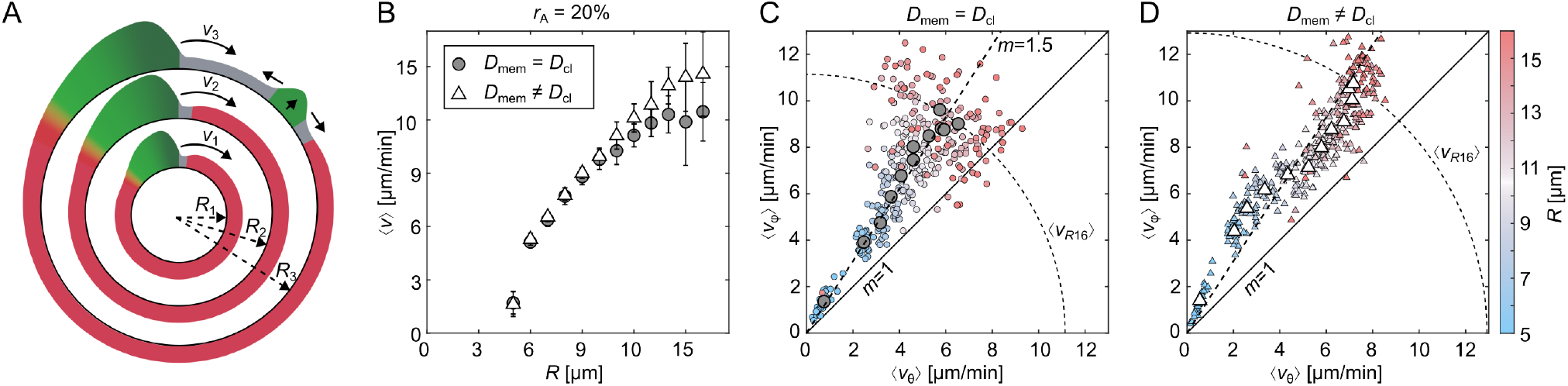
Influence of curvature and spatially limited diffusion on PIP3 waves in cells with *r*_A_ = 20%. **A** shows a scheme on the effect of restitution and dispersion in three one-dimensional rings of different radius, where *v*_2_ (*R*_2_ ) *> v*_1_ (*R*_1_ ) and *v*_3_ (*R*_3_ ) *> v*_2_ (*R*_2_ ). The average speed ⟨*v*_*R*3_ ⟩ of all waves may be smaller or larger than *v*_2_, depending on the spatiotemporal dynamics of secondary waves (upper right in *R*_3_ ). The colors indicate the excited (green), inhibitory (red) and rested states (grey). **B** shows the average of the mean velocities ⟨*v*⟩ depending on the cell radius *R* (restitution). **C** and **D** show the relation between the mean velocity components of the transversal ⟨*v*_*θ*_⟩ and longitudinal ⟨*v*_*φ*_⟩ directions for each simulation depending on *R* and the diffusion properties, respectively. The average values sorted by radius are shown by large grey circles (**C**) and large white triangles (**D**). The dashed line shown in **C** is the linear fit of the average values (large grey circles) with a slope of *m* = 1.5, and is plotted for comparison in **D** too. The dashed arc depicts the theoretical average of the mean velocities ⟨*v*_*R*16_⟩ at *R* = 16 μm.

### Membrane Curvature Restricted Diffusion

Figure 3B statistically compares the dispersion relationships between PIP3 waves on membranes (*r*_A_ = 20%) with uniform and contact line limited diffusibility, as grey circles and white triangles. The cell radius *R* defines the mean velocity ⟨*v*⟩ for both cases fairly well. For larger cells, a dispersion of ⟨*v*⟩ is observed, as secondary PIP3 waves lead to more complex wave-wave interactions. Although the dispersion relationships are very similar, there is a profound difference for the mean velocity components. Figures 3C and D show the relations between the transversal ⟨*v*_*θ*_⟩ and longitudinal ⟨*v*_*φ*_⟩ velocities of those waves for each simulation. In case of uniform diffusibility, the average velocity ratio (grey circles) for each cell size is constant at a slope of about *m* = 1.5 (dashed line). With the increase in cell size an increase of the dispersion of possible velocity ratios is observed, ranging from unity *m* = 1 up to a ratio of about *m* = 3 (see small magenta circles). Contrarily, PIP3 waves on membranes with contact line limited diffusibility have a slope beyond *m* = 1.5 for smaller cells and a slope beneath *m* = 1.5 for larger cells (Fig. 3C). Only at *R* = 9 μm a comparable aspect ratio (*m* = 1.5) is observed. Also the increase in dispersion for larger cells is visibly reduced compared to the case of uniform diffusibility. The lower velocity ratio and lower velocity dispersion for larger cells can be explained by the pinned PIP3 wave tips (singularities) on the contact line that further enables and stabilized the formation of the secondary waves on the contact line between the two wave tips (see also Fig. 2D). PIP3 waves in smaller cells are slowed down by their inhibitory wave tail, but stabilized by the cell asymmetry and guided along the contact line, which leads to an increased velocity ratio (*m >* 1.5).

### Bifurcation of Lipid Dynamics

For larger cells, PIP3 domains have the tendency to stabilize their propagation directionality either in longitudinal or transversal direction. Figures 4A and B show two example cells with *R* = 12 μm, *D*_mem_ = *D*_cl_, and *r*_A_ = 20%. The propagation direction is stable once a certain propagation direction is set after about 1 × 103 s. For cells with diffusion limited membranes this tendency gets broken and mainly transversal propagation is observed (Figs. 3C and D). This enforced pattern stabilization, can be explained by the bifurcation theory. So we hypothesize two stable solutions and quantify the velocity tuples ⟨*v*_*θ*_⟩ and ⟨*v*_*φ*_⟩ by a *k*-means unsupervised, clustering algorithm with either two solutions *k* = 2 when a significant difference between the calculated clusters is observed, or one solution *k* = 1 (no bifurcation). Figure 4C shows the result for the case of uniform diffusion on cells with *r*_A_ = 20%. The blue up- and red down-pointing triangles define the two identified clusters for cells with the same cell radius. The grey diamonds depict the cases where no bifurcation was identified, which was for either the smallest (*R* = 5 μm) or the largest simulated radius *R* = 16 μm. Cells with the largest radius show almost no cases of prominent longitudinal domain direction. Thus, ⟨*v*_r_⟩ is located within the lower branch that shows more balanced domain propagation between ⟨*v*_*θ*_⟩ and ⟨*v*_*φ*_⟩ (red down-pointing triangles). The larger symbols depict the average position of the identified clusters. Similarly, we analyzed the results for diffusion limited membranes (Fig. 4D). Here, only at the point of the slope transition (*m* ≈ 1.5, see Figs. 3C and D) for radii *R* = 9 μm and *R* = 10 μm a bifurcation is identified.

**Figure 4.**
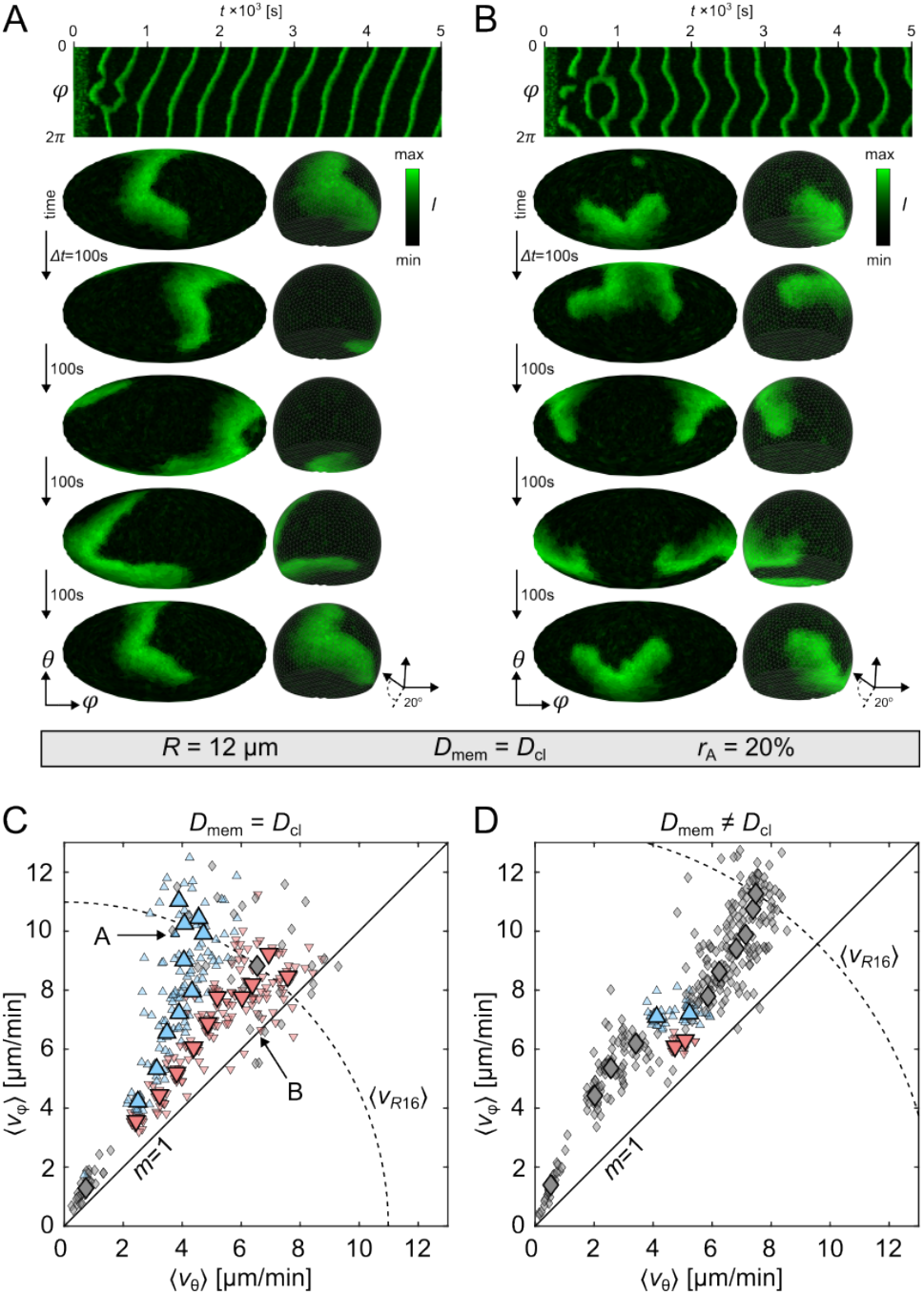
Longitudinal and transversal stable domain dynamics in cells with *r*_A_ = 20%. **A** shows longitudinal PIP3 wave dynamics with stable pinning sites on the bottom and top of the membrane. **B** shows transversal PIP3 wave dynamics with stable pinning sites around the equator of the membrane. Shape, size and diffusion are the same for both cells. The 3D views show the ventral membrane from the bottom with an inclination angle of 20°. **C** and **D** show the by *k*-means analysis sorted data for the diffusion properties *D*_mem_ = *D*_cl_ and *D*_mem_ ≠ *D*_cl_, respectively. Shown are the relation between the mean velocity components of the transversal ⟨*v*_*θ*_⟩ and longitudinal ⟨*v*_*φ*_⟩ directions for each simulation depending on *R*. Blue upwardand red downward-pointing triangles illustrate the upper and lower branch of the bifurcated data. Data shown with diamonds are not analyzed by *k*-means. The solid black line indicates the slope *m* = 1. The dashed arc depicts the theoretical average of the mean velocities ⟨*v*_*R*16_⟩ at *R* = 16 μm. The illustrated wave dynamics shown in **A** and **B** are indicated in **C** by arrows.

To proof our bifurcation hypothesis without loss of generality, we additionally performed simulations for cells with *r*_A_ = 0% and *r*_A_ = 30%, applied the same *k*-means analysis, and considered the mean velocity ratio

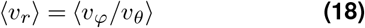

as a function of the cell radius *R*. This should not be confused with the ratio of the mean velocity components ⟨*v*_*θ*_⟩ */* ⟨*v*_*φ*_⟩ . Figure 5 shows the comparison between uniform diffusion and diffusion limited membranes. The color of the whisker boxes depict the clusters, as shown in Figs. 4C and D. Grey color depicts no bifurcation (*k* = 1) and the blue and red colors define the two clusters of the typical domain propagation patterns. Figure 5A shows the case of spherical cells (see also Fig. 2A). The median of all obtained ⟨*v*_*r*_⟩ is about unity, independent of the cell radius, as there is neither a topological asymmetry in the cell shape nor any other membrane property that may lead to a statistically reoccurring symmetry break in the pattern formation on the confined cell membranes.

**Figure 5.**
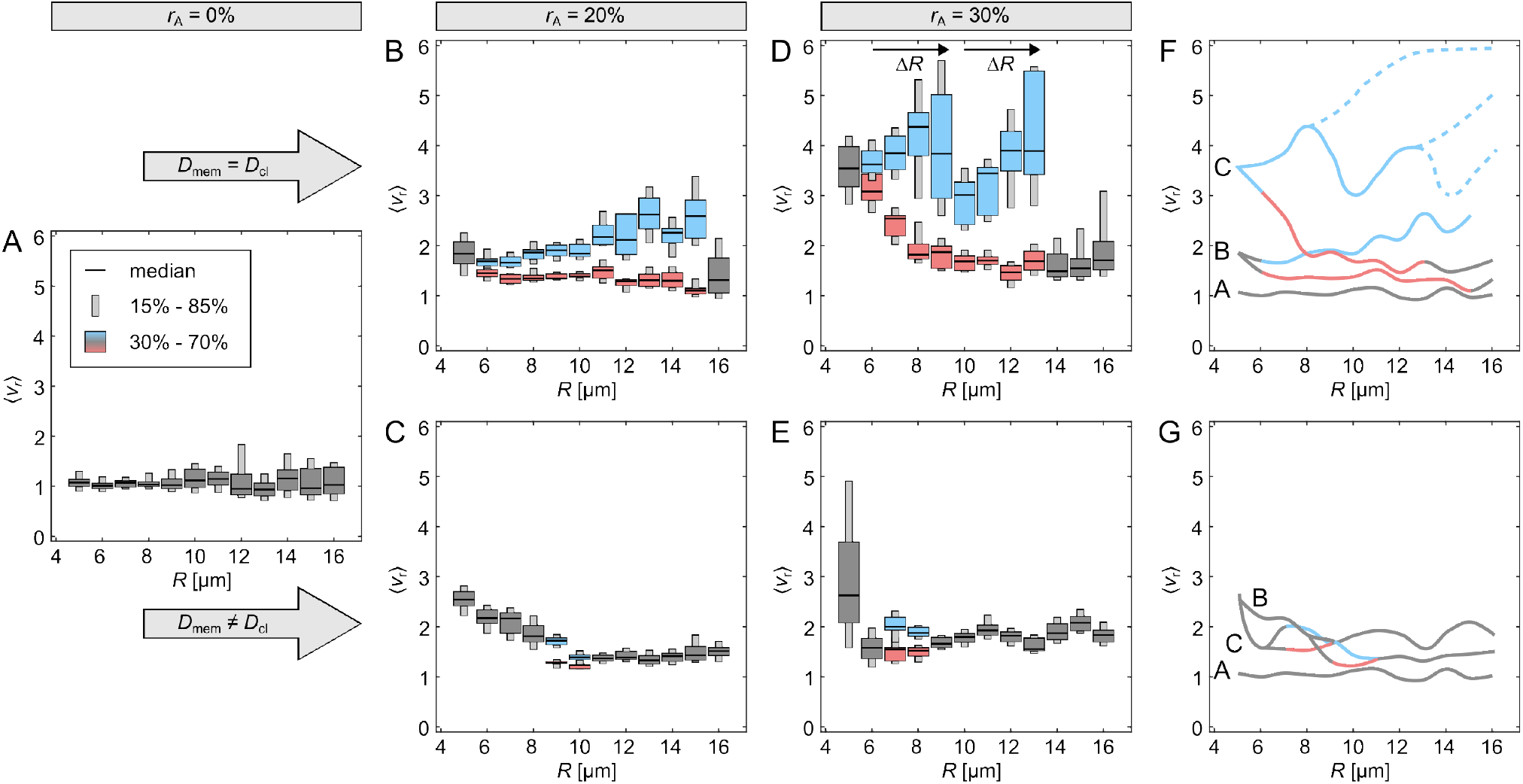
Bifurcation of the wave velocity ratio in confined membranes. **A** to **E** show whisker plots of the mean velocity ratio ⟨*v*_r_⟩ depending on the shape (*r*_A_), size (*R*) and diffusion properties of the membranes. The blue upper and red lower whiskers correspond to the by *k*-means analysis sorted relation between the mean velocity components of the transversal ⟨*v*_*θ*_⟩ and longitudinal ⟨*v*_*φ*_⟩ directions (see Fig. 4**C** and **D** for comparison). Whiskers shown with grey color are data that are not analysed by *k*-means. **F** to **G** show for comparison the smoothed median approximation of the data with uniform diffusion (*D*_mem_ = *D*_cl_) and membrane curvature reduced diffusion (*D*_mem_ ≠ *D*_cl_). Prolonged dashed lines indicate extension of the unstable hypothetically drawn bifurcation branching.

Figures 2B-E show ⟨*v*_*r*_⟩ for the combination of nonspherical cells (*r*_A_ = 20% and *r*_A_ = 30%) and different membrane diffusion properties (*D*_mem_ = *D*_cl_ and *D*mem ≠ *D*_cl_). Generally, small cells (*R* = 5 μm) show no bifurcation. With the increase in cell size ⟨*v*_*r*_⟩ decreases and eventually saturates at a low value with almost no cases of larger ⟨*v*_*r*_⟩, i.e. dominant longitudinal velocity ⟨*v*_*φ*_⟩. The second branch (blue data) shows independently of the cell topology an initial increase in ⟨*v*_*r*_⟩ when *D*_mem_ = *D*_cl_. With an increase in *R*, ⟨*v*_*r*_⟩ destabilizes until no such patterns are observed anymore. Especially for the case of *r*_A_ = 30% (Fig. 5D), a sudden drop in ⟨*v*_*r*_⟩ at *R* = 10 μm is observed, which recovers to larger ⟨*v*_*r*_⟩ until the disappearance of those patterns at *R* = 14 μm. This can be explained by an additional bifurcation with unstable solutions, as indicated by the scheme shown in Fig. 5F. The stable solutions are indicated by solid lines in the respective color. The assumed unstable solutions are indicated as blue dashed lines. It is one possible explanation, although this is a purely hypothetical solution, which we cannot back up by data. The periodic increase of the second branch (blue data) repeats at about Δ*R* = 4 μm, indicating a relationship between dispersion properties of the PIP3 domain dynamics and the confined membrane topology. One might identify a third period for *R >* 13 μm due to the occurrence of a few cases that fall in that region, which is indicated by the elongated grey whisker boxes (see also Supplementary Materials Fig. S3A). However, we refrain from defining a third period, as we cannot back up the data statistically. For the same reasons we also do not define a third lower branch for a few simulations that resulted in ⟨*v*_*r*_⟩ ≈ 1 (Fig. S3A).

The diffusion limited membranes show a different branching type (Figs. 5C and E). A reverse period-doubling bifurcation (antimonotonicity) in ⟨*v*_*r*_⟩ is observed (Dawson et al., 1992; Wouapi et al., 2019). In both cases the bifurcation is less prominent, as it only ranges for about Δ*R* = 2 μm and shows minor variations in ⟨*v*_*r*_⟩ . Arguably, differences in ⟨*v*_*r*_⟩ are weak compared to the case of *D*_mem_ = *D*_cl_, thus, they may not be bifurcations after all. However, we refer to Figs. 4D and S3B, where a clear difference in ⟨*v*_*φ*_⟩ between the two stable solution are observed. We leave it up to the reader and future investigations to argue about this findings.

### Experimental comparison

Latrunculin A-treated *Dictyostelium* cells have a typical size of about *R* = 5 μm to *R* = 6 μm. The inhibition of actin-polymerization leads to a rounding up of the cell membrane with an adhesive membrane ratio of up to about 30% (Hörning and Shibata, 2019), which is regulated by the cortical tension of the cell (Dai et al., 1999; Yang et al., 2008). Periodically transversal and longitudinal traveling PIP3 domains, as well as, spatially localized temporal PIP3 domains are observed depending on the size, shape and phenotype of the *Dictyostelium* cells (Hörning and Shibata, 2019). In numerical simulations, similar pattern formation can be observed. While larger cells exhibit traveling wave-like domains (Fig. 2), smaller cells exhibit more chaotic unstable domain dynamics (Fig. 6A). The interaction of the domains with their own inhibitory tails are stronger in smaller cells due to the limited membrane size and their confinement. Thus, the smaller the cell, the more unstable the domain dynamics.

**Figure 6.**
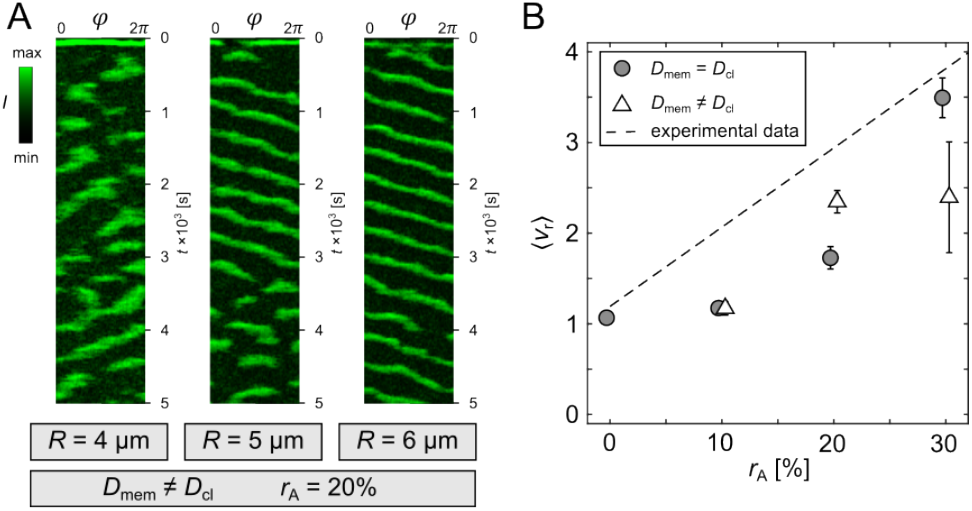
Comparison to experimental data of *Dictyostelium* cells. **A** shows three kymographs of simulations with different cell radii: chaotic (left, *R* = 4 μm), semi-stable (middle, *R* = 5 μm) and stable (right, *R* = 6 μm) PIP3 domain dynamics (*D*_mem_ ≠*D*_cl_, *r*_A_ = 20%). **B** shows the in Hörning et al. 2021 (Hörning et al., 2021) experimentally obtained analytical relation between *r*_A_ and ⟨ *v*_r_⟩ with the dashed line. The grey circles and white triangles show the comparison to the simulations with *D*_mem_ = *D*_cl_ and *D*_mem_ ≠*D*_cl_ for realistic *Dictyostelium* cell sizes of *R* = 5 μm and *R* = 6 μm only.

To validate the experimentally suggested membrane curvature influenced diffusion in *Dictyostelium* (Hörning et al., 2021; Honda et al., 2021), we compare the in Hörning et al. 2021 (Hörning et al., 2021) experimentally quantified relation between *r*_A_ and ⟨*v*_r_⟩ with the numerical results of this study (Fig. 6B). In case of uniform diffusion (*D*_mem_ = *D*_cl_, gray circles), perfectly round (*r*_A_ = 0%) and strongly adhered (*r*_A_ = 30%) cells are in good agreement with the experimental relation. However, simulations of the intermediate cell topologies (*r*_A_ = 10% and *r*_A_ = 20%) do not agree well. The simulations underestimate the experimental data, and thus, indicate a missing cell property or dynamic in the model. In the case of membrane curvature reduced diffusion (*D*_mem_ ≠*D*_cl_), the domain guiding effect of the contact line significantly increases ⟨*v*_r_⟩ for *r*_A_ = 20%, whereas there is almost no measurable effect for cells with *r*_A_ = 10%. Cells with *r*_A_ = 30% show more unstable solutions (as shown in Fig. 6A), and therefore, lead to a broader distribution of possible ⟨*v*_r_⟩ (see also Fig. 5E, *R* = 5 μm).

The results indicate that the model that considers membrane curvature reduced diffusion represents more closely the experimental data. Cells with larger adhesion ratio (*r*_A_ = 30%) follow an opposite trend, however they are barely observed, and therefore, do not account for the majority of experimentally observed cells (Hörning and Shibata, 2019; Hörning et al., 2021). Despite the improvements to the model, that more realistically represent the experimental findings compared to uniform membrane diffusion, there is still something missing that remains to be elucidated in experiments. A possible explanation might be linked to the diffusion properties of the crowded and structural cytosolic environment, that can lead to heterogeneous intracellular diffusion (Kanakubo et al., 2023) . The current model does not account for that, but considers a simplified linear global PTEN uptake only (see Eq. 17). Another experimental finding that still needs to be understood is the vertical gradient of PIP3 signaling activity observed in *Dicytostelium* cells (Hörning et al., 2021), which was obtained by spatial fluctuation analysis (Peng et al., 1994). The vertical quantification of the fluctuation parameter showed a lower value on top of the cell that increases toward the equatorial membrane with a constant slope in most cells. The cause of that general trend remains to be elucidated, and therefore, could not be considered in the current model.

## Conclusions

We investigated PIP3 domain dynamics with and without curvature limited diffusion at the contact line of the cell membrane in the first part of this study. Without loss of generality, we considered larger cells up to *R* = 16 μm, as those dynamics might be observed in other cell types. We found a Hopf-like bifurcation for the observed PIP3 dynamics in case of uniform membrane diffusion independently of the topology of non-spherical cell membranes (Fig. 5B and D). The domains propagate along either the longitudinal or transversal direction, which can be visualized with *v*_r_(*R*). In both cases, a pair of singularities (spiral wave cores) is observed. In case of longitudinally directed domain propagation, one singularity propagates along the contact line and the other one freely at the top of the cell membrane (Fig. 4A). Contrarily, for transversally directed domain propagation, both singularities are horizontally aligned and rotate as spiral waves with opposite directionality in the proximity of the equatorial height of the cell (Fig. 4B). For larger cells, only the latter is observed, as secondary PIP3 domains may appear, forcing the domains into transversal direction. A reverse period-doubling bifurcation (antimonotonicity) is observed in *v*_r_(*R*) for membranes with curvature limited diffusion at the contact line for intermediate-sized non-spherical cells (Fig. 5C and E). This type of bifurcation is observed at the transition between the initially decreasing *v*_r_(*R*) and the in *v*_(_*r*) observed asymptotic limit for larger cells. Thus, PIP3 domains in smaller cells, i.e. before this transition, are strongly influenced by the contact line, and PIP3 domains in larger cells are mainly influenced by (1) the asymmetry of the membrane topology and (2) the contact line as the main source of secondary waves (Fig. 2D). Because the influence of the contact line decreases significantly for large cells, *v*_r_ becomes comparable for cells with and without curvature limited diffusion (see Supplemental Material Fig. S4 for *R* = 16 μm). The limit of *v*_r_ depends on the strength of adhesion, which means the larger *r*_A_ the larger *v*_r_. Perfectly round cells (*r*_A_ = 0%) have a *v*_r_ ≃ 1. Contrarily, fully adhered cells with intact actin-cytoskeleton, as cultured on glass bottom dishes, have most likely a much larger *v*_r_ than the case considered in this study, i.e. *v*_r_ ≫ 2. This is because such cells have an estimated *r*_A_ ≃ 50%, as the surface-adhered membrane and the upper free membrane are of approximately the same size. Further, it would be interesting to consider membrane topologies that model tissue embedded cells and cells that form macropinosomes (Lim and Gleeson, 2011).

In the latter part of this study, we compared our results with experimental findings observed on 3D *Dictyostelium* cell membranes (Hörning et al., 2021). We restricted the comparison to cells with radii of *R* = 5 μm and *R* = 6 μm, as experimentally observed before (Hörning and Shibata, 2019). The introduced curvature limited diffusion reflects more closely the experimental data for *r*_A_ ≃ 20%. Contrarily, for less adhered cells (*r*_A_ ≃ 10%) virtually no effect has been observed. The fraction of adhered membrane is too small, hence does not affect ⟨*v*_r_⟩ . While the observed discrepancy between the experiments and numerical results might be mainly caused by the still unclear origin of the vertical gradient of PIP3 signaling activity on *Dicytostelium* cell membranes, as mentioned before, there are also other improvements to the model that can be made. The restitution and dispersion properties could be tuned to account for the biological variance in the dynamics, and the cytosolic dynamics more realistically modeled, including the effect of the intracellular crowding. The diffusion constant might differ for the adhered membranes, and the diffusion constant on the strongly curved membrane is arbitrarily chosen, because there are no experimental data available yet. Also the quantification of the PIP3-domain dynamics can be improved. Only lower scalar harmonics are considered for enabling PIP3peak detection. Larger scalar harmonic degrees increase the accuracy of peak detection, but might induce errors due to unwanted peak-jumps.

In this paper, we investigate the effect of diffusion by considering the one-dimensional curvature defined by the adhesion angle of the cell membrane only. However, a better measure might be the Gaussian curvature *K*. It is defined on the surface as the product of the two principle curvatures, as *K* = *κ*_1_*κ*_2_. On the contact line of symmetric actin-inhibited cell membranes, as considered in this study, *κ*_1_ = *κ*_*θ*_(*θ*_cl_) and *κ*_2_ = *κ*_*ϕ*_ = *R*_cl_(*ϕ*)−1 with *R*_cl_ as the horizontal radius along the contact line. In future, the model might be also applied and compared to more complex biological processes and dynamics, such as to migrating cells (Honda et al., 2021; Yang et al., 2023) and macropinocytosis (Lim and Gleeson, 2011) by adding mechno-chemical signaling pathways that control filamentous actin assembly (Beta et al., 2023).

## Acknowledgements

We want to thank Dr. Tatsuo Shibata for the discussion in the early stage of this study, the High-Performance Computing Center Stuttgart (HLRS, Stuttgart, Germany) for the support using the NEC cluster, and Judith Brock, Torsten Bullmann, and Ingrid Weiss for the fruitful discussions and suggestions to improve the manuscript.

## Author Contributions

SE and MH implemented the model and analyzed the data. MH conceptualized and designed the research and wrote the manuscript. All authors contributed to the article and approved the submitted version.

