## Supplemental Table and Figures for "Self-organization of PIP3 signaling is controlled by the confinement, topology, and curvature of the cell membrane"

**Table 1.** Reaction constants and initial concentrations of the model.

| Parameter | value | dimension | short description |
|---|---|---|---|
| [PIP2] | 700 | d.l. | initial global PIP2 concentration |
| [PIP3] | 5 | d.l. | initial global PIP3 concentration |
| [PTEN] | 2 | d.l. | initial global PTEN concentration |
| [PTEN]$_{total}$ | 0.1 | $\mu M$ | total concentration of PTEN |
| $q$ | 45 | molecules$\cdot \mu m^{-2} \cdot s^{-1}$ | PTEN-independent PIP2 supply rate |
| $V_{PI3K}$ | 500 | molecules$\cdot \mu m^{-2} \cdot s^{-1}$ | max. velocity of PIP2 phosphorylation by PI3K |
| $K_{PI3K}$ | 3500 | molecules$\cdot \mu m^{-2}$ | Michaelis constant of PI3K phosphorylation reaction |
| $V_{PTEN}$ | 15 | $s^{-1}$ | dephosphoryl. rate of PIP3 by PTEN |
| $K_{PTEN}$ | 50 | molecules$\cdot \mu m^{-2}$ | Michaelis constant of PTEN phosphorylation reaction |
| $\lambda_{PIP3}$ | 0.2 | $s^{-1}$ | PTEN-independent PIP3 degradation rate |
| $\lambda_{PIP2}$ | 0.002 | $s^{-1}$ | PI3K-independent PIP2 degradation rate |
| $\lambda_{PTEN}$ | 1.0 | $s^{-1}$ | dissociation rate of PTEN from membrane |
| $V_{ass}$ | 1300 | molecules$\cdot \mu m^{-2} \cdot \mu M^{-1} \cdot s^{-1}$ | association rate of PTEN to membrane |
| $K_{PIP2}$ | 3000 | molecules$\cdot \mu m^{-2}$ | Michaelis constant of PIP2 for PTEN associated to membrane |
| $K_{PIP3}$ | 120 | molecules$\cdot \mu m^{-2}$ | half-maximum concentration of [PIP3] |
| $\chi$ | 0.001 | $\mu M \cdot \mu m^{-2} \cdot molecules^{-1}$ | transform surface to volume concentration |

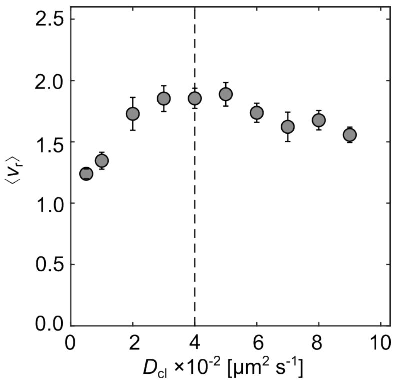


**Figure S1.** Determination of the optimal diffusion on the contact line of the membrane. The optimal diffusion $D_{\text{cl}} = 0.04$ µm$^2$/s$^{-1}$ (dashed line) was selected for cells with $R = 6$ µm, $r_{\text{A}} = 20\%$ and $D_{\text{mem}} = 0.2$ µm$^2$/s$^{-1}$. For each condition 20 simulation were computed, and the average $\langle v_{\text{r}} \rangle$ was calculated. The error bars indicate the standard error.

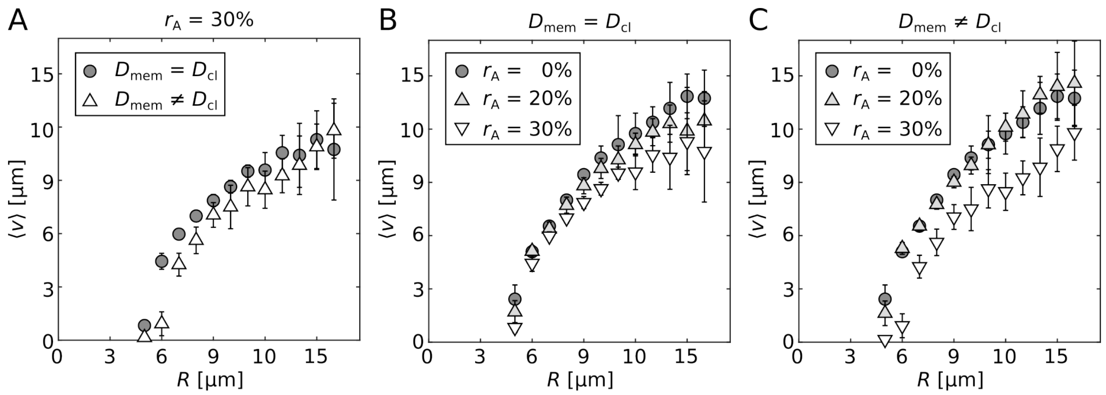


**Figure S2.** Comparison of restitution curves. **A** shows the restitution curves, i.e. the average of the mean velocities $\langle v \rangle$ depending on the cell radius $R$ for $r_A = 30\%$. **B** and **C** show the restitution curves of three cell shapes ($r_A$) for $D_{mem} = D_{cl}$ and $D_{mem} \neq D_{cl}$, respectively.

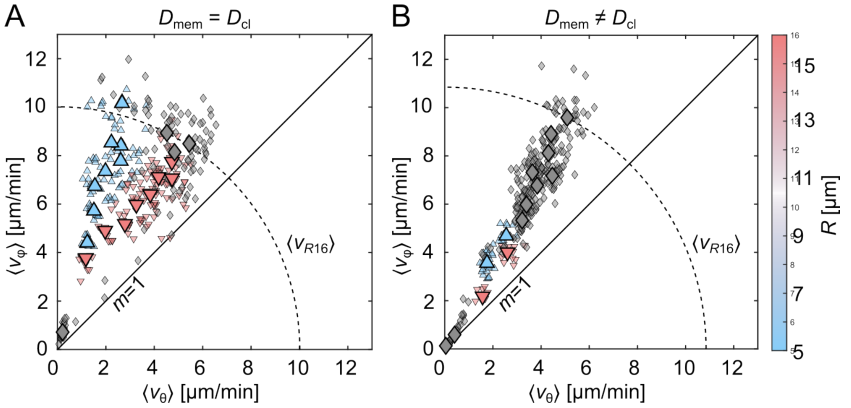


**Figure S3.** Longitudinal and transversal stable domain dynamics in cells with $r_A = 30\%$. **A** and **B** show the by $k$-means analysis sorted data for the diffusion properties $D_{mem} = D_{cl}$ and $D_{mem} \neq D_{cl}$, respectively. Shown are the relation between the mean velocity components of the transversal $\langle v_\theta \rangle$ and longitudinal $\langle v_\varphi \rangle$ directions for each simulation depending on $R$. Blue upward- and red downward-pointing triangles illustrate the upper and lower branch of the bifurcated data. Data shown with diamonds are not analyzed by $k$-means. The solid line indicates the slope $m = 1$. The dashed arc depicts the theoretical average of the mean velocities $\langle v_{R16} \rangle$ at $R = 16$ µm.

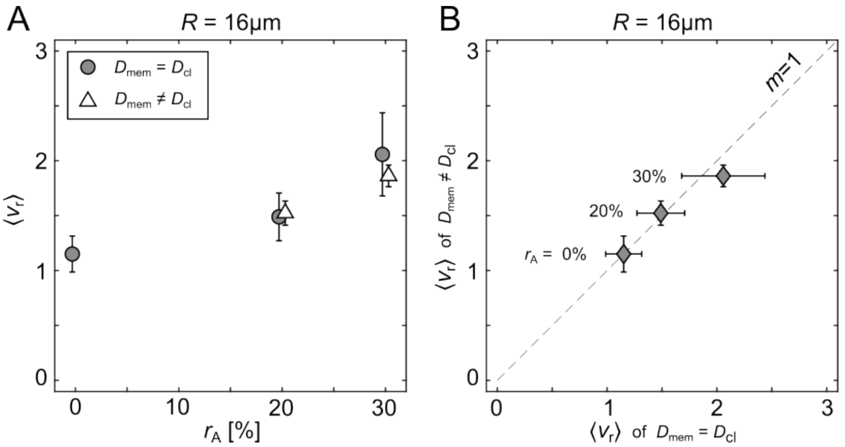


**Figure S4.** Comparison of domain dynamics between diffusion properties for $R = 16$ µm. **A** shows the relation between $r_A$ and $\langle v_r \rangle$ for the simulations with $D_{mem} = D_{cl}$ (gray circles) and $D_{mem} \neq D_{cl}$ (white triangles). **B** shows the $\langle v_r \rangle$ of $D_{mem} = D_{cl}$ and $D_{mem} \neq D_{cl}$ plotted against each other for different $r_A$. The dashed line shows the slope of unity.
